# A novel behavioral apparatus for spontaneous exploration and operant conditioning of social information under spatial conditions in rats

**DOI:** 10.1101/2024.10.02.616150

**Authors:** Taylor B. Wise, Victoria L. Templer, Rebecca D. Burwell

## Abstract

**Background:** Evidence from the fields of evolutionary biology and neuroscience supports the theory that spatial cognition and social cognition share neural mechanisms. Rodent models are widely used to study either spatial or social cognition, but few studies have explored the interactions between the spatial and social cognitive domains due to the lack of appropriate paradigms.

**New method:** Our study introduces the Vertical Maze (VM), a novel behavioral apparatus designed to measure multiple aspects of spatial and social cognition. The VM features a standard 3-chamber maze positioned above multilevel columns allowing for the presentation of conspecifics at varying spatial distances and familiarity levels. This arrangement enables conspecifics to serve as discriminative stimuli for both social and spatial spontaneous and goal-oriented assessments. The three-dimensional design of the VM allows rats to use multisensory cues to judge distance, direction, and social identity of conspecifics.

**Results:** In the present study, we found that rats can 1) discriminate the spatial distance of conspecifics located below them in an operant conditioning task, and 2) discriminate social novelty when conspecifics are presented below at the near, middle, and far distances in a spontaneous exploration task. Critically, it was necessary for rats to explore all levels of the maze to perform these discriminations.

**Comparison with existing methods:** This new method advances the field by permitting the presentation of social information (conspecifics) at different spatial distances. The use of conspecifics to serve as stimuli for both social and spatial discriminations allows more direct comparison of behavioral measures across these information domains. Importantly, the presentation of conspecifics as stimuli below the 3-chamber level of the maze engages auditory, visual, and olfactory systems, encouraging a robust multisensory representation of conspecifics presented at a distance.

**Conclusions:** Our results confirm that the VM is an effective tool for studying both spatial and social cognition, facilitating the development of novel automated tasks in these areas. This new method opens new avenues for investigating the neural and cognitive foundations of spatial and social behavior, as well as for exploring the possibility of shared mechanisms across these cognitive domains.

## INTRODUCTION

Growing evidence from the fields of evolutionary biology and neuroscience supports the theory of shared spatial and social cognitive mechanisms (Parkinson & Wheatley, 2013; Schafer & Schiller, 2018; Tavares et al., 2015). Few studies, however, have addressed possible interactions between these two cognitive domains, in part due to experimental differences in how spatial and social cognition are measured. Here, we review tasks used for studying spatial and social behavior in rodents, and then propose a new approach that is optimized for studying how information is processed in and across these cognitive domains.

Rodents provide a useful animal model for studying social behavior. Most studies in rodents focus on spontaneous and innate social behaviors, such as social dominance, recognition of conspecifics, preference for social novelty, and motivation for social interaction (Figure 1). Commonly used tasks for evaluating spontaneous social behavior include the 3-chamber sociability and social novelty test (Figure 1A) (Acikgoz et al., 2022; Crawley, 2004; Moy et al., 2004), tests for social preference/avoidance (Berton et al., 2006; Haller & Bakos, 2002; Toth & Neumann, 2013), the tube dominance task {Harris, 2021 #103;Ishaq, 2023 #104;Lindzey, 1961 #568;Wilson, 1968 #569}, and the spontaneous social interaction test (File & Hyde, 1978; File & Seth, 2003). The 3-chamber test places animals in an apparatus consisting of three chambers where they can explore holding cages that are either empty or occupied by a familiar or novel conspecific (Figure 1A). Normal rats and mice prefer to explore conspecifics more than empty cages and novel conspecifics more than familiar ones (Moy et al., 2004; Templer et al., 2018). Similar tests can be accomplished with the Social Y-maze, an apparatus in which two of the arms end with cubic wire holding cages for presenting conspecifics (Weber-Stadlbauer et al., 2017). While the 3-Chamber test is a time-efficient method for assessing social behavior and social recognition memory, there are limitations. For example, results could be ambiguous because of exploratory bias, and the method does not allow for extended training or continuous measures of social behaviors.

**Figure 1.**
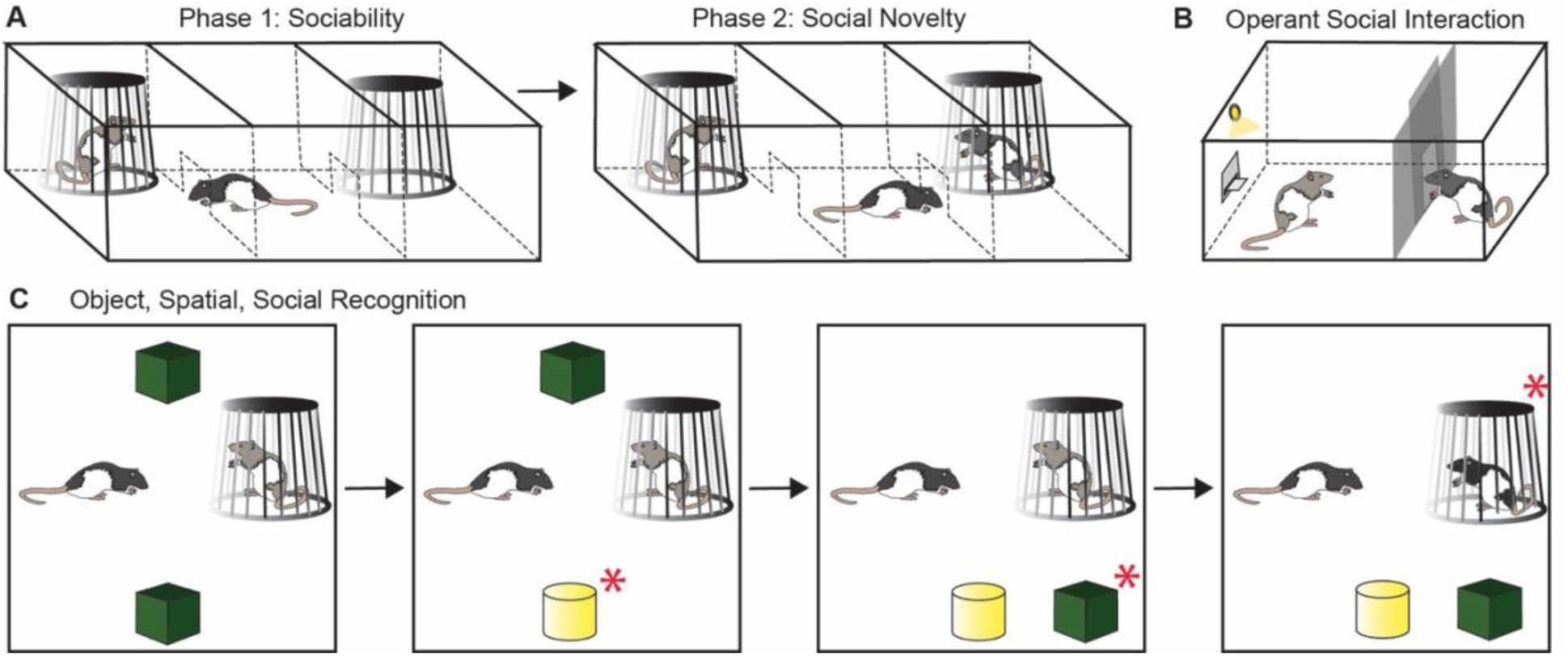
Standard tests of social interaction. A) Three-Chamber Sociability and Social Novelty Test. This test assesses general sociability and interest in social novelty in rodent models. Rodents prefer to spend time with a conspecific over an empty cage (Sociability), and they will normally prefer a novel conspecific over a familiar one (Social Novelty). B) Operant conditioning paradigm for social interaction. Social interaction is rewarding, and rodents will learn to lever press for social interaction with a conspecific. This paradigm quantifies motivation for social experience. C) Object, Spatial, Social Recognition task (adapted from Lian et al., 2018). The red asterisk indicates the predicted preferences for each phase of the task.

To address the limitations of spontaneous exploration tasks, some investigators have used operant conditioning approaches in which rodents learn operant responses to obtain access to a conspecific as a reward (Figure 1B) (Martin & Iceberg, 2015; Trezza et al., 2011). The specific methods vary, but generally involve the rodents performing an action (e.g. lever press or nose poke) to gain access to brief social contact with a conspecific. In one operant social self-administration task, mice learned to lever press for access to brief social interaction with a same sex cage mate (S. S. Lee et al., 2024; Ramsey et al., 2023). In a similar operant model, rats learned to lever press for access to brief social interactions with another rat even as interactions progressively required more effort (Achterberg et al., 2016). These approaches allowed for studying volitional social reward-seeking under operant conditions. A variation on this task used an apparatus with three chambers separated by one-way doors (Borland et al., 2017). Rodents could press levers to gain access to either a chamber containing a conspecific or an empty chamber. Operant paradigms can also be used for higher order associations with a social component. In a “social incentivization of future choice” task, mice were trained to nose poke for two equally preferred food rewards (Hu et al., 2022). One food was then paired with social interaction. In probe trials mice preferred the nose poke response associated with the socially paired food reward. Rats can also be tested for prosocial behavior in operant tasks. In an automated two-choice task, rats could choose between levers to both earn a reward and provide a reward for a partner rat (Kentrop et al., 2020). Rats showed a preference for the lever that rewarded both rats. Overall, operant conditioning approaches provide more quantitative and reliable measures of social behavior, longer testing periods, and more elaborate experimental designs. They provide more control over the animal’s behavior and attention and can be designed to test a variety of aspects of social behavior.

Rodents are also skilled navigators and have historically played a prominent role in spatial cognition research. Classic tests for rodent spatial cognition include navigational tasks like the Morris Water Maze (Morris, 1981), the Barnes Maze (Barnes, 1979), and the Olton radial arm maze (Walker & Olton, 1979) as well as Y-maze and T-maze tests. The water maze evaluates spatial learning and memory by requiring rodents to find a hidden platform submerged in a circular pool of opaque water. Rodents must use distal visual cues to navigate and locate the platform. It is particularly useful for studying spatial navigation abilities. The Barnes maze is a dry-land maze consisting of a circular platform with multiple holes around the perimeter, one of which leads to an escape box. Rodents must use spatial cues to find the escape hole, making this test suitable for assessing navigation, spatial learning, and memory with less stress compared to the water maze. The radial arm maze consists of a central hub with multiple arms radiating outwards from the center. Each arm may contain a food reward. It can be used to study both working memory (remembering which arms have been visited within a session) and reference memory (remembering which arms are consistently rewarded across sessions). This maze is useful for studying decision-making and the effects of various treatments on memory. The T-Maze and Y-Maze are also used to investigate spatial working memory and decision-making. In spontaneous alternation tasks, rodents must remember which arm they previously visited and choose the alternate arm on subsequent trials. These mazes are also used for reward-based tasks to study spatial learning and memory. Other approaches to investigating spatial cognition include spatial recognition tasks like object exploration (Warburton & Brown, 2015) and place preference (Bardo et al., 1989). Spatial recognition tasks require initial encoding of a spatial environment and then detection of any environmental changes that occurred. To construct specific spatial environments, these tests commonly use objects that can be altered in terms of location, identity, or configuration (Blaser & Heyser, 2015; Dix & Aggleton, 1999; Save et al., 1992; Warburton & Brown, 2015). Like social preference tests, rats exhibit increased exploration of novel spatial changes, which can be used as a measure of recognition memory.

Despite rodents’ capabilities for social and spatial cognition, very few studies have broached the topic of how these cognitive domains may intersect. Of the available studies, results are limited to how conspecific behavior influences subjects’ spatial exploration. In one observational learning study, rats that were allowed to first observe Barnes Maze navigation by demonstrator rats solved the task by finding the escape hole significantly faster than their demonstrators (Yamada & Sakurai, 2018). Rats are also capable of learning a rewarded spatial location based solely on observing a conspecific find the food reward (Doublet et al., 2022). When tasked with learning multiple locations of food rewards in a matrix of possible locations, rats with prior knowledge of the rewarded locations were not impacted by a conspecific partner, but rats without prior knowledge of the rewarded location were strongly influenced by social cues of a conspecific that did have prior knowledge (Bisbing et al., 2015). Another study compared how three factors – access to food, the presence of a partner, and past spatial experience of the food location – influenced collection of food in a spatial environment. Exploring with a conspecific had the most influence in spatial decision-making and spatial location knowledge had the least (Dorfman & Eilam, 2018). In an open field exploration task, solo rats explored more because rats in pairs spent the majority of exploration sessions interacting socially rather than spatially (S. L. T. Lee et al., 2023). This suggests that – like other types of stimuli that are prioritized only when useful – social information modulates behavior under specific conditions.

Studies like the ones discussed above inform our understanding of social cognition, spatial cognition, and social influence over spatial behavior. These studies do not, however, address the potential behavioral and mechanistic similarities between spatial and social information processing. In one notable exception, recognition memory of spatial and social information was investigated in a single paradigm (Lian et al., 2018). In this task, rats were presented with two objects and a conspecific in each exploration session. Over a series of sessions, objects changed in identity and spatial location, and the conspecific was replaced with a different animal (Figure 1C). Rats were equally able to detect changes in spatial and social information. This study permitted evaluation of how object identity, object location, and conspecific identity might compete for an animal’s attention and motivation to explore. However, while rats in the study preferred to spend more time with conspecifics as compared to objects, such preferences might result in experimental confounds while attempting to understand how spatial and social information processing interact, such as unequal interest in inanimate-object vs live-animal stimuli.

One solution to the potential confounds induced by the strong preferences for social interaction in rodents would be to use conspecifics as stimuli for non-social tests. The use of conspecifics for both spatial and social discriminations could provide new opportunities for addressing open questions about spatial-social domain interplay in rodent models. Here, we introduce the Vertical Maze (VM), a novel apparatus capable of capturing spontaneous social preferences and goal-driven decision-making under social or spatial demands (2A). Features of the apparatus include 3 testing chambers positioned above two vertical columns in which social stimuli can be positioned at three spatial distances from the subject. The 3 testing chambers closely resemble a 3-chamber testing apparatus with additional features to allow for operant conditioning. Automated hardware in the two outer chambers of the VM include levers, lights, pellet dispensers, and guillotine doors. Testing chamber floors are interchangeable. The floors can be solid to provide a simple 3-chamber apparatus, permeable to allow subjects to see, smell, and hear stimuli in the lower levels, or equipped with passageways to permit subjects to explore and spatially map the vertical columns below the three-chamber level. In two experiments we confirmed rats will explore and can identify conspecifics presented below the testing chamber at three spatial distances. We also confirmed rats can learn operant tasks in the maze when stimuli are conspecifics presented below the testing chambers in the vertical columns.

In Experiment 1, we conducted a social preference test that measured spontaneous exploration of various social contexts. The design was adapted from the 3-chamber test but presented conspecifics below the testing chambers. This experiment addressed several practical questions. First, what conditions are necessary for rats to attend to stimuli presented below the floors of testing chambers? Attention may depend on prior mapping of space below the testing chambers. To address this question, some subjects received additional habituation sessions prior to testing in which they traversed vertical space in the columns below the test level (top) of the maze (2B). Second, is social information presented in the vertical columns sufficient for judgement of relative familiarity? If demonstrator rats are farther away, is more time needed for them to become familiar to the subject? In the first phase of the 3-chamber test (Sociability), it is important that subjects have sufficient experience with the demonstrator so that there is a difference in relative novelty between the original and novel demonstrator in second phase (Social Novelty) (Figure 2C). Otherwise, demonstrators in Social Novelty might produce competing novelty preferences for the subject that confound results. In other words, the original demonstrator may still be perceived by the subject as somewhat novel and thus diminish preference for the second, truly novel demonstrator. To address this concern, we first conducted either a standard 10-minute or extended 20-minute Sociability test. Extended time during Sociability allowed for twice the amount of time subjects could become familiar with the original demonstrator before the introduction of a novel demonstrator in Social Novelty. Performance during Social Novelty was then compared across the standard and extended duration groups. Potential impacts on familiarization were examined for all spatial distances. For our third question – does distance from the testing chamber impact social preference? To address this question, we examined sociability and social novelty preferences when conspecifics were presented below the testing chamber at near, middle, and far distances from the subject (Figure 2D).

**Figure 2.**
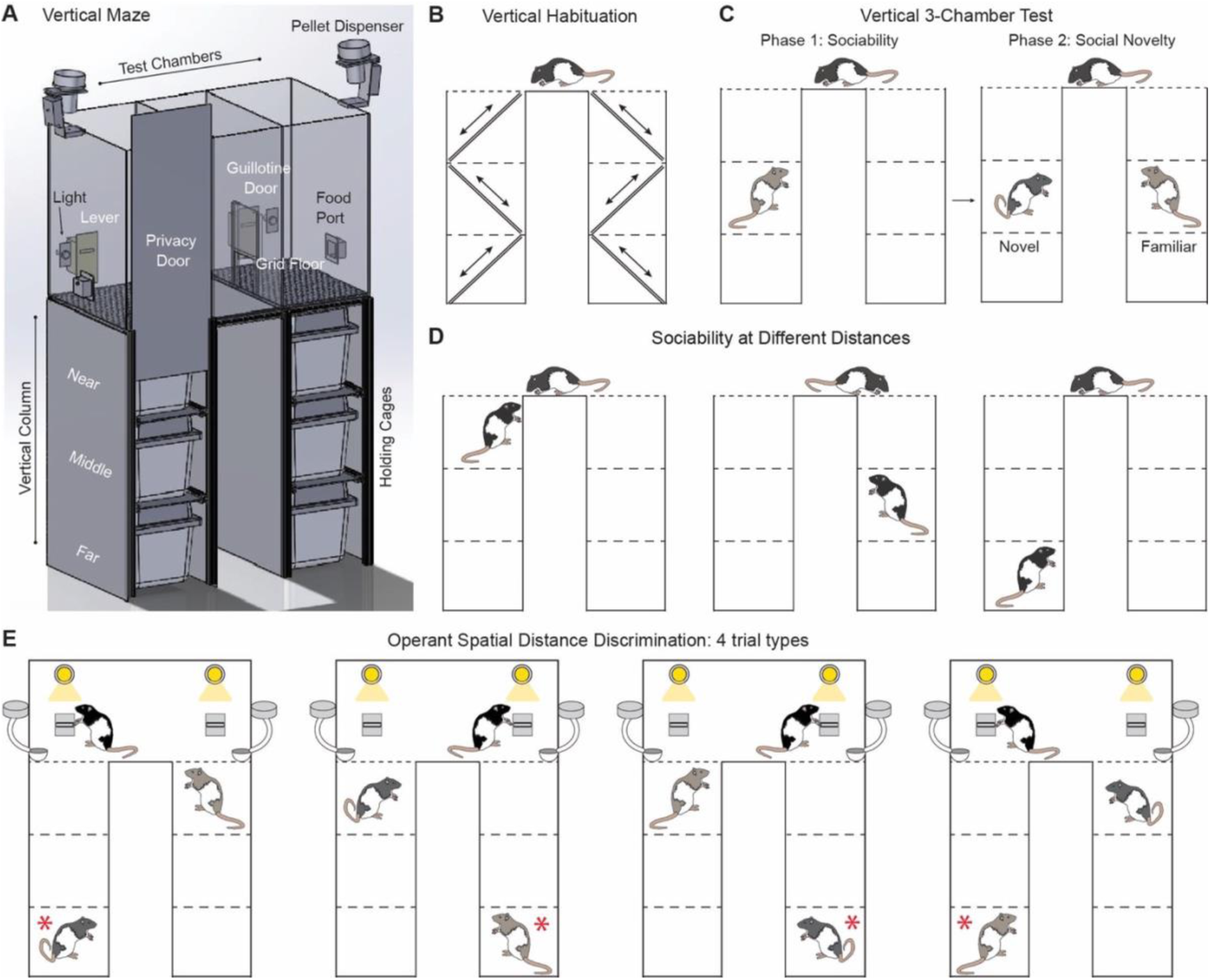
Vertical Maze (VM) and example behavioral paradigms. A) Rendering of the VM. Major components include 3 testing chambers that the subject can explore and two vertical columns where social stimuli (conspecifics in holding cages) can be positioned at three spatial distances from the subject. Automated hardware on either side of the VM includes levers, lights, pellet dispensers, food ports, and guillotine doors. The testing chamber floors are interchangeable and can be, for example, opaque, translucent, or grid-like. B) Schematic of the habituation procedure for allowing rats to explore vertical columns below the testing chambers of the VM. C) Sociability and social novelty testing in the VM. The sociability phase includes a demonstrator and an empty holding cage to measure rats’ preference between social and non-social contexts. The social novelty phase includes the demonstrator from the prior phase (considered familiar) as well as a novel demonstrator to measure social novelty preference. Conspecifics can be presented at three spatial distances from the subject. D) Sociability testing at three possible spatial distances. A conspecific was presented at the near, middle, or far distance. Spatial distance of conspecifics in Sociability remained the same for Social Novelty. E) Schematic of the Vertical 3-Chamber Spatial Distance Discrimination Operant task showing four trial types. Rats were trained to press a lever corresponding to either the near or far demonstrator for a food reward. Individual demonstrator rats were presented equally often in the near and far, and east and west location, such that the discrimination could be only based on spatial distance. The red asterisk shows the correct choice in the far-trained condition.

In Experiment 2, we conducted an operant conditioning test that measured goal-oriented discrimination between two spatial distances (Figure 2E). Conspecifics were used as stimuli and presented below testing chambers at the far vs. near spatial distance. Choice was defined by pressing the lever in the testing chamber positioned above either distance. This experiment addressed whether the VM can be used for operant learning of spatial information while using social stimuli. Across experiments, our results shed light on how spatial and social information processing may interact during multiple behavioral paradigms.

We found that the VM can be used to study sociability and social novelty preferences when conspecifics are presented below the maze, even at the farthest level. However, it was necessary to provide subjects with exploratory sessions of the vertical columns prior to testing. More exploration time for subjects in the sociability phase, however, was not necessary for normal behavior in the social novelty phase. Finally, subjects were able to learn an operantly conditioned distance discrimination in the maze. Results from this study are among the first to report rat’s spatial and social behavior in a shared paradigm. The development of the VM will allow new approaches to the study of spatial and social information processing in rodent models, including addressing questions about their neural bases and shared mechanisms.

## MATERIALS AND METHODS

### Subjects

Eighty six adult, male Long Evans rats were used in this study (Charles River Laboratories, Boston, MA, USA). Experiment 1 included 54 subjects and 26 non-subject conspecifics used as demonstrators. Social interaction between subjects and demonstrators never occurred outside of testing. Experiment 2 included an additional 6 subjects that were trained on an operant task. Four of these rats had undergone sham surgeries in which they were anesthetized and given craniotomies and allowed to recover at least two weeks prior to behavioral training. Rats were pair-housed in ventilated cages. Rats were put on food schedules to maintain 85-90% of free feeding weight. Access to water was unrestricted. Rats were placed on a 12:12 reversed light-dark cycle so that testing was conducted at a consistent time during the dark phase. Prior to any testing, rats were well-handled by experimenters. All procedures were conducted according to the NIH guidelines for the care and use of rats in research and approved by the Institutional Animal Care and Use Committee (IACUC) at Brown University.

### Apparatus

The VM is a modular apparatus capable of testing numerous parameters including sociability, social novelty preference, social discrimination, spatial discrimination, and a variety of operant discriminations (Figure 2A). The VM comprises a three-chamber testing platform with solid Plexiglass walls (91.4 x 52 x 61 cm) positioned above two vertical columns (33 x 52 x 87.6 cm, each). The two outer (testing) chambers accommodate interchangeable floors made of perforated aluminum, wire grid, or a plexiglass floor that includes passages for exploring the vertical columns. Flooring in the middle (start) chamber was always solid plexiglass. Hardware in each testing chamber included an automated lever, light, speaker, pellet dispenser, and pellet receptacle (Med Associates, Inc.), as well as a custom guillotine door to control subject access to adjacent chambers. Two vertical columns positioned below the two outside chambers were designed to accommodate holding cages at three vertical levels such that the floors of the three levels were 29.2 cm, 58.4 cm, and 87.6 cm below the floor of the 3-chamber testing arena. Vertical columns were constructed of T-slotted aluminum framing and corrugated plastic sheets with sliding privacy doors to create a dark, quiet environment for demonstrators. The plastic holding cages were equipped with wire bar lids. To permit exploration of the entire apparatus, including the vertical columns, holding cages with access holes were placed at the three vertical levels and the top holding cages included access to the 3-chamber maze. Holding cages with access holes were used only for a vertical exploration phase prior to testing. Control of all hardware and behavioral tracking was conducted via Ethovision XT (Noldus Information Technology). A high-speed video camera was positioned above the VM and provided real-time feedback to tracking software for behavior-dependent hardware control.

One advantage of the VM as compared to the 3-chamber maze is that the holding cages are likely less stressful for the demonstrators. Standard holding cages are 20×40 cm in size (Harvard Apparatus) and do not include privacy from the larger apparatus. The VM holding cages are larger and can be equipped with bedding, access to food and water, and enrichment items. Moreover, they are enclosed in the dark, private vertical columns. Given that stress signals are transmitted between individuals (Fernández-Vargas et al., 2022; Johnston, 2003; Portfors, 2007) and can have lasting impacts on behavior (Morozov, 2018; van der Kooij & Sandi, 2012), demonstrator comfort is an important consideration. Demonstrator stress levels during social tests are not often documented despite potential impacts on subject performance. The VM demonstrator holding cages may allow longer test trials and ensure against often undetected stress-related behavioral confounds.

### Behavioral Procedures

#### Habituation

Before testing, subjects and demonstrators were habituated to the VM. There were two habituation conditions for subjects. In one, subjects were habituated only to the top level, and in the other subjects were given additional habituation sessions in which they also explored the vertical columns. In the initial habituation, subjects were first placed in the start chamber with guillotine doors open and food pellets were scattered throughout to promote exploration. Perforated aluminum flooring in the outer chambers ensured that subjects could not physically access the vertical columns during habituation. Subjects were habituated for 20 minutes a day for two days. Some subjects were given additional habituation sessions in which they had access to one testing chamber at a time along with the holding cages below (Figure 2B). Food rewards were scattered throughout. Criteria for completing vertical habituation (VH) was travel from the bottom holding cage to the testing chamber on each side of the maze, which occurred in 2 or 3 sessions of up to 30 minutes per session. Demonstrators were never present for subject habituation. Demonstrators were separately habituated by being placed in their assigned holding cages in either the west or east vertical column for 20 minutes a day for two days. To promote a comfortable environment, holding cages contained bedding, enrichment toys, and food pellets, and the privacy doors were closed. Automatic hardware in the testing chambers was sporadically run during demonstrators’ habituation.

#### Vertical 3-Chamber Sociability and Social Novelty (V3CSSN) Task

The V3CSSN task was used to address questions in Experiment 1. The task consists of two phases: Sociability and Social Novelty. In the sociability phase, a conspecific is presented below the floor on one end of the maze and no conspecific is presented on the other side. We predict that normal rats will spend more time in the chamber over the conspecific than in the chamber with no conspecific. In the social novelty phase, subjects are presented with the same conspecific on one side of the maze and a novel conspecific on the other side. We predict that normal rats will spend more time in the chamber with the novel conspecific. To examine whether prior exploration of the entire VM was required for processing information presented below the testing chambers, we compared performance of control subjects and subjects that had also undergone vertical habituation (VH) (Figure 2B). The control group did not undergo any vertical habituation.

Social preference testing procedures were adapted from the 3-Chamber sociability and social novelty task (Crawley, 2004; Moy et al., 2004) with the exception that demonstrators were presented at different spatial distances below the subject (Figure 2C). Wire grid flooring in the west and east chambers allowed subjects to perceive demonstrators without physical contact. In the sociability phase, control subjects were presented with a novel demonstrator at only the nearest spatial distance in either the west or east vertical column (Figure 2C, left). VH subjects were presented with a novel demonstrator placed at the near, middle, or far spatial distance in either the west or east vertical column. The opposite vertical column remained empty. Demonstrator side was counterbalanced across control and VH subjects. Demonstrator distance was counterbalanced for VH subjects. To initiate the trial, a subject was placed in the center chamber and guillotine doors were lifted to allow for exploration of both the west and east testing chambers.

To evaluate whether spatial distance can impact the pace of familiarization with a novel demonstrator, we manipulated exploration time in the sociability phase of the V3CSSN test. During Sociability, control and VH – Standard subjects were allowed to explore for 10 minutes, whereas VH – Extended subjects were allowed to explore for 20 minutes. Social Novelty exploration time was 10 minutes for all groups. At the end of Sociability, all subjects were placed back in the center chamber and guillotine doors were closed for an intertrial interval (ITI) of two minutes. In the social novelty phase, a second, novel demonstrator was placed in the previously empty column at the same vertical level as the demonstrator from before (now familiar to the subject) (Figure 2C, right). Spatial distance remained consistent across phases. To initiate the trial, doors were again lifted, and subjects explored testing chambers for 10 minutes. At the end of testing all rats were returned to their home cages. Testing chambers and vertical columns were cleaned and aired out with a handheld fan to reduce odor transfer among subjects and demonstrators.

#### Vertical 3-Chamber Spatial Distance Discrimination Operant (V3CSDO) Task

Prior to testing subjects were habituated to the VM testing chambers identical to habituation in Experiment 1 with the exception that all subjects were also given vertical habituation. Following habituation, rats were shaped to press a lever for a food reward in the west and east testing chambers. Chamber side was alternated each session for all subjects. Subjects began with magazine training in which a pellet was dispensed into the receptacle every 30 seconds for 15 minutes, and the lever was not present. After two sessions of magazine training, rats were hand-shaped to press the lever. Rats were then moved to autoshaping in which they were required to lever press for a reward with no assistance from the experimenter. Autoshaping was complete when a rat pressed the lever 10 times in a single session for each side of the maze.

In the V3CSDO task, subjects learned to discriminate demonstrators based on spatial distance (Figure 2E). In each trial, one demonstrator was placed at the top level of one vertical column and another demonstrator rat was placed at the bottom level of the opposite vertical column. Half of the subjects were in the far condition (rewarded for choosing the farthest demonstrator) and half were in the near condition (rewarded for choosing the nearest demonstrator) for the duration of spatial training. For each subject, the same two equally unfamiliar demonstrators were used in each trial for the duration of training. Demonstrator position was pseudo-randomly organized such that each demonstrator was positioned at the top and bottom and appeared in both the west and east columns for an equal number of trials per session, as shown in Figure 2E. Counter-balancing distance and side ensured that spatial distance was the only relevant dimension for a correct choice.

Spatial distance training consisted of 20 trials per session for 12 daily sessions. Before each trial subjects were placed in the closed center chamber and demonstrators were positioned at the near and far spatial distances. At the start of each trial, guillotine doors were lifted, lights were turned on, and levers were extended in the testing chambers. Each chamber included a wire grid floor that allowed subjects to perceive demonstrators positioned below. Choice was indicated by a lever press in either the west or east chamber. Correct choices were signaled by a brief tone, and three pellets were dispensed before lights were turned off and levers were retracted. Each session began with five discovery trials. If a subject’s first choice was incorrect, the pressed lever was retracted, and the correct lever remained extended until pressed, resulting in the tone and delivery of 3 pellets. Those trials were scored as incorrect. After 5 discovery trials, an incorrect choice terminated the trial with no reward. At the end of each trial, subjects were placed back in the center chamber with doors closed and demonstrators were positioned for the next trial. Demonstrators were assigned individual holding cages to limit potential cross-odor contamination. Testing chambers and vertical columns were cleaned and aired with a handheld fan between subjects to reduce odor cues. Subjects received 12 daily sessions.

### Analysis

Sociability and social novelty preference was defined as exploration of the west or east testing chamber in which the demonstrator was positioned underneath. Preference for each phase was analyzed using a discrimination ratio (DR) which quantifies the difference in exploration as a proportion of the total exploration time (Ennaceur & Delacour, 1988). In Sociability, a positive DR indicated a preference for the social context over the side of the maze in which there was no demonstrator, while in Social Novelty a positive DR indicated a preference for the side of the maze with the novel conspecific over the side with the familiar conspecific. In both cases, a DR significantly above zero indicated a preference for sociability or social novelty.

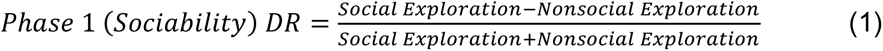

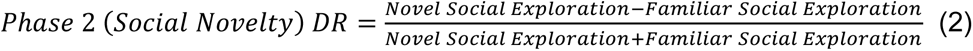

Consistent with prior studies utilizing DR analyses (Heimer-McGinn et al., 2017), data acquired in the first three minutes of each phase was analyzed to provide an accurate depiction of rats’ initial preferences before behavior was extinguished. One-sided, one-sample *t*-tests were used to determine whether DRs were significantly different than zero. A significant *t*-test indicated sociability or social novelty preferences. Depending on the question, DRs were analyzed by one-way analysis of variance (ANOVA) with group as the between-subject variable or a repeated measures analysis of variance (RMANOVA) with group as the between-subject variable and phase (sociability vs. social novelty) as the within-subject variable. Outlier analyses were included in Experiment 1 with outliers beyond two standard deviations from the mean treated as missing data and replaced with the closest data point within the normal distribution (Blaine, 2018). In Experiment 2, accuracy and latency data were analyzed by RMANOVA with trained correct distance as the between-subject variable and session as the within-subject variable. Behavioral measures for both experiments were collected using Ethovision XT and analyzed using IBM SPSS Statistics (Version 29).

## RESULTS

### Experiment 1: Sociability and social novelty in the VM

The control group, which did not undergo vertical habituation and was trained only at the near distance, did not show a preference for the column with the conspecific in the Sociability test or the novel conspecific in the Social Novelty test, whereas the VH group did show the predicted preferences (Figure 3A). In the comparison of the DRs to zero, the VH subgroup trained at the near distance showed significant Sociability (t_7_ = 16.000, p < 0.000) and Social Novelty (t_7_ = 2.929, p < 0.022) preference, whereas controls showed no preference in Sociability (t_7_ = −0.582, p > 0.579) and a familiarity preference in Social Novelty (t_7_ = −2.496, p < 0.021, Table S1, left two columns). The VH group, including all distance conditions, significantly preferred Sociability (t_23_ = 11.219, p < 0.000) and Social Novelty (t_23_ = 3.488, p < 0.002). For Sociability, the DRs were significantly above zero for all spatial distances (p-values ranged from 0.0001 to 0.001, Table S1, middle left column). For Social Novelty, the DRs were significantly above zero for the near (p < 0.011) and far (p < 0.007) distances, but not the middle distance (p > 0.181), Table S1, middle left column). RMANOVA of the DRs confirmed that the control and VH near distance subgroup were significantly different for Sociability (F_1,15_ = 14.169, p < 0.002) and Social Novelty (F_1,15_ = 14.522, p < 0.002) at the near spatial distance (Table S2). A significant group difference was also found when comparing controls to the VH group including all distance conditions for Sociability (F_1,31_ = 29.086, p < 0.000) and Social Novelty (F_1,31_ = 12.310, p < 0.001).

**Figure 3.**
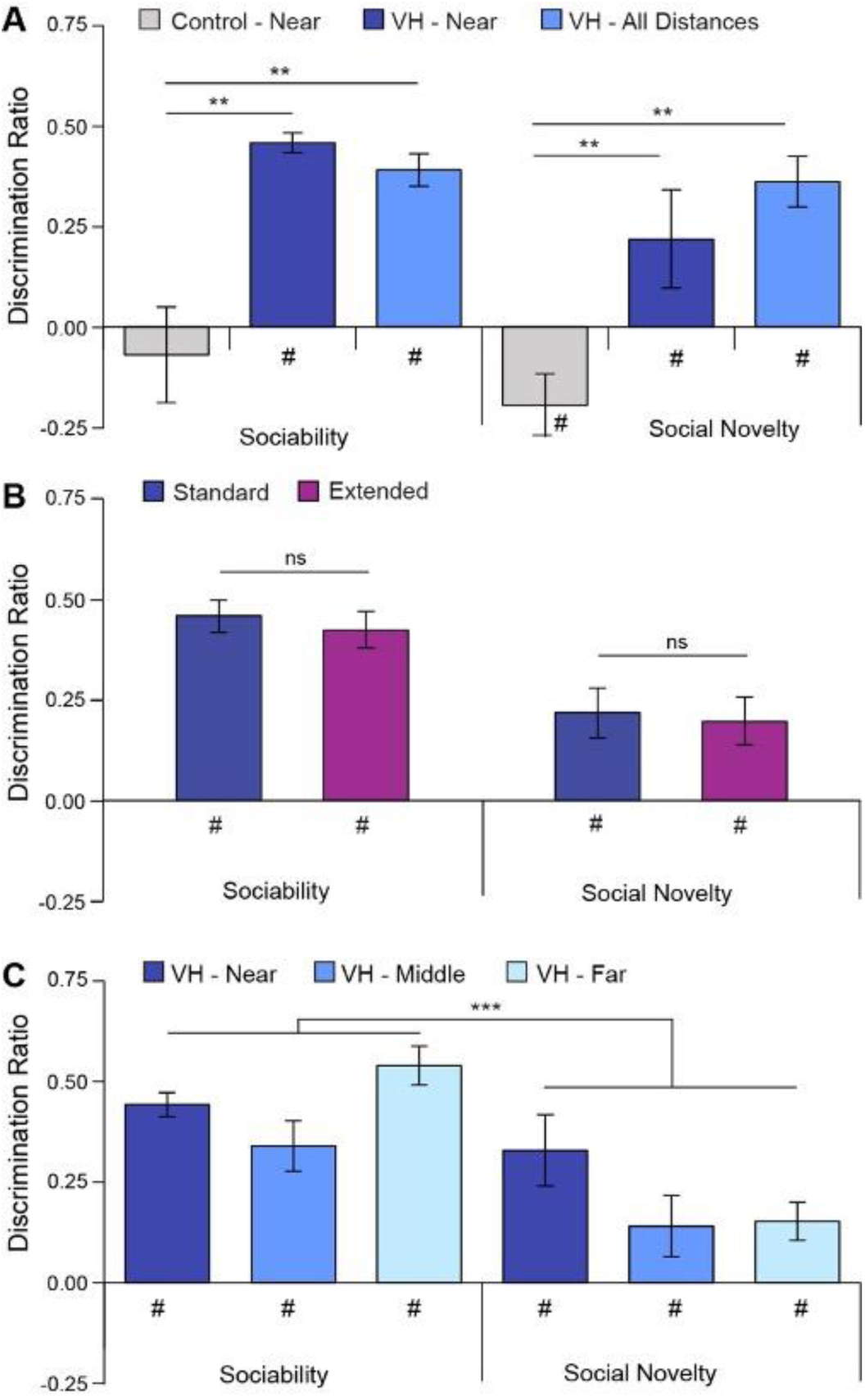
Discrimination ratios for Sociability and Social Novelty tests under different conditions. A) Impact of vertical habituation (VH) on Sociability and Social Novelty. Control rats were trained only at the near distance and VH rats were trained in the near, middle, or far distance. Experimental rats, but not control rats, were given pre-test exploration sessions in which they traversed vertical space in the east and west columns. For Sociability, the Control - Near group preferred the conspecific side of the maze significantly less than the VH - Near group or the VH – All Distances group. For Social Novelty, the VH groups preferred the novel conspecific whereas controls did not. B) Impact of exploration duration on sociability and social novelty preferences in the VM. The standard group explored for 10 minutes during Sociability while the extended group explored for 20 minutes. Both groups received VH prior to testing. Both groups preferred the conspecific during Sociability and the novel conspecific during Social Novelty. C) Sociability and social novelty preference for groups of VH rats trained at the near, middle, and far spatial distances. All three distance groups showed DRs significantly above zero for both Sociability and Social Novelty, DRs were significantly different between phases with stronger preferences for Sociability as compared to Social Novelty. *** p < 0.000; # below bars indicates DR scores significantly different from zero.

We were interested in whether subjects might need more exploration time in the VM apparatus during Sociability to become sufficiently familiar with the demonstrator to allow a preference for the new demonstrator during Social Novelty. Overall, the DRs for the VH-extended group were similar to the VH-Standard group (Table S1, middle two columns). For Sociability, the DRs were significantly above zero for all spatial distances (p-values ranged from 0.0001 to 0.010). For Social Novelty, the DRs were significantly above zero for the near (p < 0.037) and marginally significantly above zero for the middle (p < 0.063) and far distances (p < 0.074). RMANOVA of the DRs for standard vs. extended exploration conditions in the sociability test showed a main effect of spatial distance (F_2,13_ = 7.007, p < 0.004), no main effect of exploration duration (F_2,13_ = 0.120, p > 0.735), and a marginal ‘spatial distance x exploration duration’ interaction (F_2,13_ = 3.325, p < 0.053; Figure 3B; Table S2). For Social Novelty, there was no main effect of spatial distance (F_2,13_ = 1.834, p > 0.180) or exploration duration (F_1,13_ = 0.017, p > 0.899), and no ‘spatial distance x exploration duration’ interaction (F_2,13_ = 0.090, p > 0.915; Figure 3B; Table S2). Thus, the standard sociability exploration time of 10 minutes appears to be sufficient to allow the preference for the novel conspecific in the social novelty test.

Because RMANOVA indicated no differences in DR between the standard and extended exploration durations, those data were combined for further spatial distance analyses. Overall, subjects showed the predicted preferences. One-sided, one-sample *t*-tests indicated that DRs were significantly above zero for all three spatial distances for Sociability (p < 0.000) and Social Novelty (p-values ranged from 0.001 to 0.046, Table S1, right column). Discrimination ratios were higher during Sociability than Social Novelty, and exploration differed across spatial distances (Figure 3C). This was confirmed by RMANOVA, which indicated a main effect of phase (F_1,28_ = 18.033, p < 0.000) and spatial distance (F_2,28_ = 4.094, p < 0.022), but no ‘phase x spatial distance’ interaction (F_2,28_ = 2.601, p < 0.083; Figure 3C; Table S2).

### Experiment 2: Operant spatial distance discrimination in the VM

During shaping, each rat successfully learned to lever press in the VM in five to 10 days with an average of 6.1 ± SE days. Following two magazine-only sessions, magazine + shaping sessions were conducted with an average of 4.9 ± SE sessions required to meet criterion. For autoshaping, average number of sessions to criterion was 1.10 ± SE. Two rats met lever pressing criterion on the final magazine + shaping session and did not require autoshaping.

In spatial distance training both the near and the far distance groups demonstrated improved accuracy across trials (Figure 4A). Early in training the near group performed better, in the middle of training the far group performed better, and by the end of training the groups were nearly identical. This was confirmed by a main effect on accuracy of session (F_11,44=_ 4.819, p < 0.001) and a significant ‘session x distance’ interaction (F_11,44_ = 2.742, p < 0.009), but no main effect of distance (p < 0.904). Latency to make a choice decreased with training for both distance groups (Figure 4B). This was confirmed by a main effect of session (F_11,44_ = 8.366, p < 0.001), but no main effect of distance (p > 0.994) or ‘session x distance’ interaction (p > 0.996). Together, these results suggest that rats effectively learned to discriminate between spatial distance of two conspecifics in an operant conditioning paradigm whether the correct choice was the near conspecific or the far conspecific.

**Figure 4.**
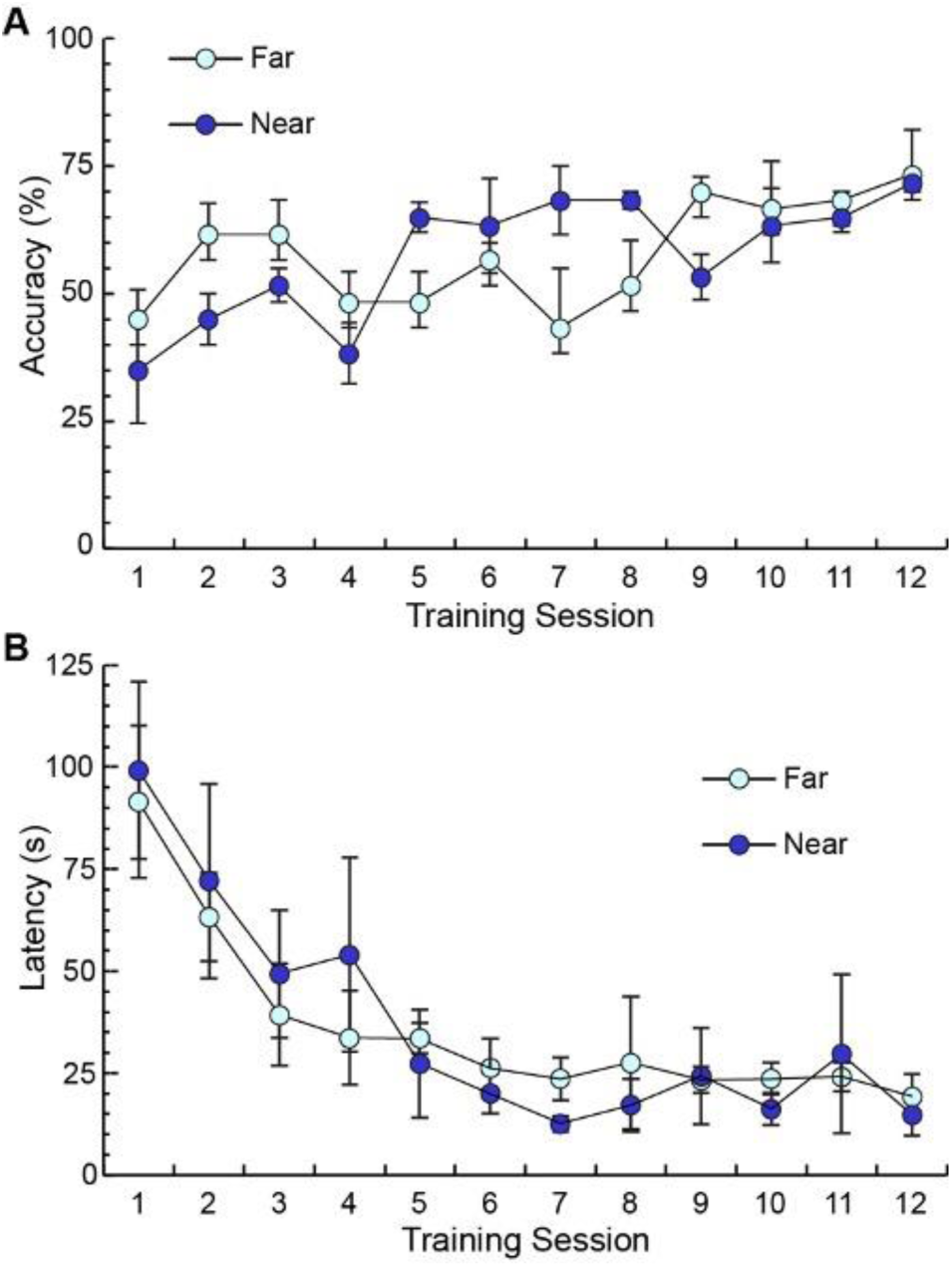
Operant conditioning of a spatial distance discrimination with conspecifics as stimuli. A) Accuracy improved with training for both near- and far-trained distance groups as confirmed by a main effect of session (p < 0.001). Although there were accuracy differences in both directions early in training, both groups were performing equally well by the end of training. B) Likewise, latency to choice decreased with training for both distance groups as confirmed by a main effect of session (p < 0.001)

## DISCUSSION

In this study we introduce the Vertical Maze, a novel and versatile apparatus that can be used for both spontaneous exploration tasks, for example to test sociability and social novelty preferences, and operant conditioning tasks, in which rats are trained to make a discrimination and are rewarded for pressing a lever. Notably, the VM permits presentation of social information, i.e. conspecifics, at different spatial distances. This allows conspecifics to serve as stimuli for both social and spatial discriminations, enabling direct comparison of behavioral measures. This innovation opens new avenues for investigating the neural and cognitive foundations of spatial and social behavior, as well as exploring the connections and parallels between these cognitive domains.

In the present study we addressed several practical questions about the feasibility of using conspecifics as stimuli in the VM. Experiment 1 demonstrated the VM’s effectiveness as a method for studying social recognition. Our results replicate sociability and social novelty preferences found in the standard 3-chamber sociability test (Moy et al., 2004; Templer et al., 2018). Moreover, DRs were significantly above zero for the near, middle, and far spatial distances indicating that rodents can recognize conspecifics at a distance. To our knowledge, this is the first reported social test to manipulate a spatial distance variable.

We also showed that exploration of vertical space below the testing floor of the VM was necessary for subjects to demonstrate sociability and social novelty preferences. Prior research shows that, for rats, direct exploration appears to improve the accuracy and stability of 3D spatial representations (Grieves et al., 2020; Grieves & Jeffery, 2017; Hayman et al., 2011; Jedidi-Ayoub et al., 2021). Thus, it could be that spatial mapping of the vertical columns is necessary for rats to attend to the spaces below the top level. In a multilevel object recognition task, rats were also habituated to all levels before training began, although the investigators did not test whether habituation was necessary for good performance (Cole et al., 2019). In our study, the control group, which had not undergone pretesting vertical habituation, displayed no preference for sociability or social novelty when demonstrators were at the near distance level, whereas the vertical habituation group showed significant discrimination at all three distance levels. This strongly suggests that exploration of the vertical columns in the VM is necessary for attending to social information presented underneath subjects.

In social assessments, rodents typically have very close access to conspecific demonstrators, thus it was possible that spatial distance might hamper social recognition. For this reason, we designed our apparatus and task to engage auditory, visual, and olfactory systems to encourage a robust multisensory representation of conspecifics presented at a distance. Indeed, social cues are communicated through all three of these systems (de la Zerda et al., 2020; Ebbesen & Froemke, 2021; Fernández-Vargas et al., 2022; Seffer et al., 2014). We found that subjects were able to identify familiarity and novelty of conspecifics presented at the near, middle, and far spatial distances. Another concern, however, was whether more exploration would be needed during Sociability such that demonstrators at greater distances were considered sufficiently familiar before Social Novelty. It might have been the case that more time was needed for subjects to become familiar with demonstrators in the far condition. We tested this possibility and found that task exploration time had no impact on discrimination for either Sociability or Social Novelty (Figure 3B). This indicates that spatial distance did not impair social familiarization in the VM and that the 10 min exploration time typically used in the 3-chamber test is also sufficient when the conspecific is presented at a distance. Discrimination ratios were lower for Social Novelty, but this is consistent with findings in the standard 3-chamber maze (McKibben et al., 2014). Together, these results validate the VM as an appropriate test for social preference when spatial distance is a variable.

The operant spatial distance discrimination task used in Experiment 2 provided evidence that rats can also distinguish the distance at which conspecifics are experienced. With training, accuracy improved and latency to choice decreased for both subjects trained on the near distance and the far distance, and by the end of training both groups were performing equally well. This opens the door for developing tasks that address whether spatial distance and social identity, otherwise referred to as social distance, are processed similarly in the brain. Given that rats can discriminate spatial distance and social identity at different distances in the VM, it is now possible to test hypotheses about the neural bases of information processing in the spatial and social information domains using either spontaneous exploration or operant paradigms. For example, it would be possible to test whether a group of rats trained on a distance discrimination transfers spatial distance information to the social domain when given a distance probe test.

## CONCLUSION

This study introduces the Vertical Maze (VM), a novel and versatile apparatus designed for both spontaneous exploration and operant conditioning tasks in rats. The VM allows researchers to 1) test sociability and social novelty preferences through spontaneous exploration tasks, 2) conduct operant conditioning tasks in which rats learn discriminations and are rewarded for lever pressing, and 3) present conspecifics at varying spatial distances, enabling their use as stimuli for both social and spatial discriminations. This innovative design facilitates direct comparisons of behavioral measures across social and spatial domains. The VM opens up new possibilities for investigating the neural and evolutionary bases of spatial and social cognition and behavior, as well as exploring potential connections between these cognitive domains.

## ACKNOWLEDGEMENTS

We would like to thank John Murphy for his assistance in engineering and constructing the Vertical Maze. We would also like to thank Jasmine Lee, Isabella Pilkington, Maya Mazumder, and Laura Betances for their assistance in data collection and Devon Poeta, MS. for her advice.

## SUPPLEMENTARY MATERIALS

**Table S1.** Impact of spatial distance on sociability and social novelty preferences.

|  |  | Control |  |  | Vertical Habituation (Standard) |  |  | Vertical Habituation (Extended) |  |  | Vertical Habituation (Standard & Extended) |  |  |
| --- | --- | --- | --- | --- | --- | --- | --- | --- | --- | --- | --- | --- | --- |
|  |  | t | df | p | t | df | p | t | df | p | t | df | p |
| Sociability | Near | -0.582 | 7 | 0.289 | 16.000 | 7 | **0.000** | 9.400 | 6 | **0.000** | 14.561 | 14 | **0.000** |
|  | Middle | - | - | - | 4.451 | 7 | **0.001** | 3.145 | 6 | **0.010** | 5.408 | 14 | **0.000** |
|  | Far | - | - | - | 14.754 | 7 | **0.000** | 6.389 | 7 | **0.000** | 11.152 | 15 | **0.000** |
| Social Novelty | Near | -2.496 | 7 | **\*0.021** | 2.929 | 7 | **0.011** | 2.170 | 6 | **0.037** | 3.731 | 14 | **0.001** |
|  | Middle | - | - | - | 0.976 | 7 | 0.181 | 1.778 | 6 | 0.063 | 1.810 | 14 | **0.046** |
|  | Far | - | - | - | 3.290 | 7 | **0.007** | 1.629 | 7 | 0.074 | 3.205 | 15 | **0.003** |

One-sided, one-sample t-tests were used to determine whether DR scores were significantly above zero, thus indicating preference for the conspecific in the Sociability test and the novel conspecific in the Social Novelty test. \*Note that the control group showed a preference for familiarity in the Social Novelty test.

**Table S2.** Impact of vertical habituation, exploration duration, and test phase on Sociability and Social Novelty.

ONE-WAY ANOVA: Impact of vertical habituation at near distance
|  | Control (n=8) |  | Vertical Habituation (n=8) |  | Control vs. VH |  |
| --- | --- | --- | --- | --- | --- | --- |
|  | Mean | StdErr | Mean | StdErr | F | p |
| Sociability | -0.070 | 0.120 | 0.390 | 0.069 | 14.169 | **0.002** |
| Social Novelty | -0.195 | 0.078 | 0.360 | 0.123 | 14.522 | **0.002** |

REPEATED MEASURES ANOVA: Impact of exploration duration and distance
|  | Standard (n=24) vs. Extended (n=21) Duration |  |  |  |  |  |
| --- | --- | --- | --- | --- | --- | --- |
|  | Within Subject |  | Between Subject |  | Interaction |  |
|  | F | p | F | p | F | p |
| Sociability | 7.007 | **0.004** | 0.120 | 0.735 | 3.325 | 0.052 |
| Social Novelty | 1.843 | 0.180 | 0.017 | 0.899 | 0.090 | 0.915 |

REPEATED MEASURES ANOVA: Strength of sociability vs social novelty
|  | Sociability vs. Social Novelty (n=45) |  |  |  |  |  |
| --- | --- | --- | --- | --- | --- | --- |
|  | Within Subject |  | Between Subject |  | Interaction |  |
|  | F | p | F | p | F | p |
| Both Phases | 4.094 | **0.022** | 18.033 | **0.000** | 2.601 | 0.083 |

A one-way ANOVA (top) was used to compare controls, who received no vertical habituation (Vh), to the VH group at the near spatial distance. Both groups received the standard exploration duration in Sociability. An RMANOVA (middle) was used to compare the standard duration group to the extended duration group at each spatial distance. Both groups received VH prior to testing. The within subject variable was spatial distance and the between subject variable was exploration duration. No group difference in exploration duration was found for either phase, therefore subjects were combined to compare strength of sociability to social novelty preference in a final RMANOVA analysis (bottom). The within subject variable was phase and the between subject variable was spatial distance.

